# Pericystic brain transcriptomics reveals molecular signatures of immune activation and neurovascular remodelling in viable and post-treatment porcine neurocysticercosis

**DOI:** 10.64898/2026.06.26.734379

**Authors:** Carla A. Apaza-Quiroz, Cielo C. Rojas-Portocarrero, Sneider A. Gutierrez Guarnizo, Emily K. Ponce-Nakatahara, Javier A. Bustos, Gianfranco Arroyo, Robert H. Gilman, Héctor H. García, Mirko Zimic

## Abstract

Neurocysticercosis (NCC), the infection of the central nervous system by *Taenia solium* larvae, is a leading cause of acquired epilepsy in endemic regions. While viable cysticerci can persist asymptomatically for extended periods, their spontaneous or drug-induced degradation triggers marked perilesional inflammation and severe neurological symptoms. Despite well-documented histopathological characterisation of these lesion states, the host transcriptional programmes associated with viable parasite persistence and early post-treatment lesion disruption remain poorly understood. To address this gap, we performed the first bulk RNA sequencing of pericystic brain tissue using a physiologically relevant porcine model of NCC. Comparing uninfected controls (n = 3), infected untreated pigs with intact viable cysts (n = 6), and antiparasitic-treated pigs with disrupted cysts (n = 3), we identified distinct transcriptional signatures associated with each disease state. Viable infection was associated with broad transcriptional changes (461 upregulated and 175 downregulated genes), characterised by local immune activation alongside suppression of blood-brain barrier (BBB) remodelling, vascular, and neuronal signalling molecular signatures. The post-treatment state with confirmed BBB disruption was associated with a smaller but directionally distinct response (160 upregulated and 57 downregulated genes), marked by inflammatory signalling and increased expression of genes associated with endothelial activation, vascular regulation, and BBB-associated remodelling. Together, these findings suggest that, while immune engagement is a feature shared across both lesion states, the BBB-associated transcriptional axis shifts substantially following treatment. These results provide an exploratory transcriptomic framework for understanding parasite persistence, treatment-induced neuroinflammation, and neurovascular remodelling in NCC, and highlight candidate pathways and genes for future mechanistic investigation.

**Author Summary:** Neurocysticercosis is a major cause of epilepsy in regions where *Taenia solium* is endemic. Brain cysts can remain viable for long periods with limited symptoms, but parasite degeneration, whether spontaneous or drug-induced, can trigger damaging neuroinflammation. In this study, we used RNA sequencing in a pig model that closely resembles human disease to characterise how brain tissue responds to viable cysts and to early treatment-induced cyst disruption. We found that viable infection was associated with local immune activation alongside reduced expression of genes involved in blood-brain barrier function. Following antiparasitic treatment, disrupted lesions showed an increased expression of genes linked to vascular and barrier remodelling. These findings suggest that the host transcriptional environment changes substantially after parasite disruption, and highlight molecular pathways that may contribute to neuroinflammation, blood-brain barrier changes, and neurological disease in NCC. As an exploratory first transcriptomic survey in this model, these results provide a candidate framework for future studies aimed at identifying biomarkers and adjunctive therapeutic targets in NCC.

## Introduction

Neurocysticercosis (NCC) is the most prevalent parasitic infection of the human central nervous system and remains a major cause of acquired epilepsy in endemic settings across Latin America, Africa and Asia (1). The disease arises when humans ingest eggs of *Taenia solium*, the pork tapeworm, through faecal-oral contamination. Released oncospheres penetrate the intestinal wall, enter the bloodstream, and develop as cysticerci preferentially in the brain parenchyma (2). Once established, these cysticerci may persist for prolonged periods, sometimes years, with minimal surrounding inflammation. This is not merely passive tolerance, as preliminary data from humans and naturally infected pigs indicate that viable parasites actively engage host immune suppression, characterised by elevated expression of anti-inflammatory mediators including IL-10 and TGF-β (2,3). Additional immune evasion strategies attributed to *T. solium* cysticerci include protection by local barriers such as the blood-brain barrier or the hemato-ocular barrier, as well as the degradation of host immunoglobulins or masking with host immunoglobulins to evade immune surveillance (4). Despite the clinical importance of these mechanisms, the host transcriptional landscape supporting parasite persistence in brain tissue has not previously been characterised by unbiased, transcriptome-wide profiling.

When cysticerci begin to degenerate, either spontaneously or after antiparasitic therapy, they elicit marked perilesional inflammation that drives many of the clinical manifestations of NCC, including seizures, headaches, hydrocephalus and focal neurological symptoms (2,3). Degenerating lesions are characterised by lymphocytic and myeloid infiltration, granulomatous tissue remodelling, and increased inflammatory mediator production, and are commonly associated with oedema and contrast enhancement on neuroimaging (2,3,5–8). Although inflammation typically resolves as lesions progress to calcification, residual scarring can perpetuate morbidity. Defining the molecular programmes that govern lesion state transitions is therefore central to understanding NCC pathogenesis and improving patient management.

Current treatment of parenchymal NCC relies on cysticidal therapy with albendazole and praziquantel, frequently accompanied by corticosteroids to mitigate treatment-associated inflammation (5). However, a major drawback of antiparasitic treatment is that parasite injury can precipitate acute host inflammatory responses as cyst antigens are released into the surrounding tissue (5,6). In the porcine model, Cangalaya et al. (6) documented this process within days of treatment initiation, when lesions show increased contrast enhancement, Evans Blue extravasation, and more intense perilesional inflammation. This is consistent with previous clinical observations in humans that perilesional oedema accompanies the onset of anthelmintic therapy, sometimes precipitating seizures or intracranial hypertension (9). Accordingly, the dominant source of NCC morbidity is the host immune response to dying parasites rather than direct mechanical damage by the cyst itself.

Because access to human brain tissue is extremely limited, much of what is known about NCC immunopathology has come from animal models. Rodent systems have been valuable for studying selected aspects of central nervous system inflammation and barrier biology, but they do not reproduce the natural host-parasite biology of *T. solium* infection (10). By contrast, pigs are the natural intermediate host of *T. solium* and constitute a physiologically relevant model for human NCC (10). Both naturally and experimentally intracarotid-infected pigs exhibit viable, degenerating, and calcified cyst stages, as well as the characteristic pattern of minimal inflammation around live cysticerci and robust granulomatous responses around degenerating ones, which parallel those observed in humans (5,10). These properties make the porcine model appropriate for studying host-parasite interactions in NCC.

Despite the clinical importance of these lesion states and the strengths of the porcine NCC model, the molecular pathways operating in lesion-adjacent brain tissue during viable infection and after antiparasitic treatment remain poorly defined. Although previous studies have used RT-qPCR panels to examine gene expression during host response (5,11), whole-transcriptome profiling has not yet been performed in any NCC animal model. This gap limits our ability to understand the mechanisms by which viable parasites maintain immunological tolerance, how treatment disrupts this balance, and which molecular pathways contribute to the ensuing neuroinflammation.

To address this, we performed the first bulk RNA-seq analysis of pericystic brain tissue from experimentally infected pigs, defining host transcriptional responses to viable infection and to early antiparasitic treatment. Given the exploratory nature of this dataset, particularly the small sample size and the absence of orthogonal validation, all findings are presented as hypothesis-generating observations rather than mechanistic conclusions. Our results identify three principal transcriptional patterns: (i) local immune engagement is a shared feature of both viable-infection and post-treatment pericystic tissue; (ii) MAPK signalling shows divergent enrichment between conditions in a manner potentially relevant to NCC epileptogenesis; and (iii) BBB-associated and vascular remodelling programmes display the most pronounced directional shift, being suppressed during viable infection and activated following treatment. These patterns provide a candidate transcriptomic framework for future mechanistic and biomarker studies in NCC.

## Methods

### Ethics statement

Samples used in this study were obtained from a previously approved pre-clinical experiment conducted under the authorization of the Institutional Animal Care and Use Committee Universidad Peruana Cayetano Heredia (approval number R-046-11-24). All procedures involving animals and tissue collection were performed in compliance with the guidelines of the Association for Assessment and Accreditation of Laboratory Animal Care and the U.S National Institutes of Health (AAALAC/NIH).

### Sample collection and study design

Pigs were allocated into three experimental groups: (i) untreated uninfected controls (n = 3); (ii) infected untreated pigs (n = 9); and (iii) infected pigs treated with a combined antiparasitic regimen of albendazole (ABZ, 15 mg/kg for five days) plus praziquantel (PZQ, 75 mg/kg for one day), and euthanised 120 hours after treatment onset (n = 6).

Brain tissue specimens used in this study were obtained from a previously established experimental porcine model of neurocysticercosis (12). Briefly, *Sus scrofa domesticus* individuals were experimentally infected through intracarotid injection of 5000 activated *T. solium* oncospheres at two months of age, and infection with viable brain cysticerci was confirmed five months post-infection by magnetic resonance imaging (MRI).

Two hours before euthanasia, pigs were intravenously perfused with a 2% Evans Blue solution (in 0.85% NaCl) at a total dose of 80 mg/kg as previously described (13). Evans Blue binds to serum albumin and is normally retained within the intact vasculature; when the BBB is disrupted, the dye extravasates and produces visible blue staining in the perilesional tissue. Cyst capsules were classified as ‘Clear’ when no dye extravasation was observed (indicating intact BBB) or ‘Blue’ when Evans Blue staining was present (indicating local BBB disruption).

Immediately after euthanasia, the brain was perfused with 2,000 mL of chilled heparinised saline (0.85% NaCl, 10 IU/mL heparin), removed from the skull, and sectioned into 10 mm coronal slices to identify cysticerci. Cysticerci with pericystic capsules were immediately transferred to RNALater for transcriptomic preservation.

For this study, pericystic tissue from Clear-capsule cysts in infected untreated pigs and from Blue-capsule cysts in antiparasitic-treated pigs was selected for RNA sequencing. Parenchymal brain tissue from uninfected pigs provided controls (n = 3). Evans Blue classification was applied to all infected animals prior to tissue selection. Of the nine infected untreated pigs, six bore exclusively Clear-capsule cysts and were retained for the infection-effect comparison (n = 6); the remaining three, which presented Blue-capsule cysts indicating BBB disruption, were excluded to avoid confounding the infection-effect contrast with barrier disruption. Of the six infected treated pigs, three presented Blue-capsule cysts and were retained for the treatment-effect comparison (n = 3); the remaining three, which presented Clear-capsule cysts, were excluded as their cysts had not achieved the expected post-treatment BBB disruption. This design (**Fig 1**) avoids the confounding that would arise from pooling cysts of mixed BBB integrity within groups, and ensures that each contrast reflects a biologically homogeneous tissue source.

**Fig 1.**
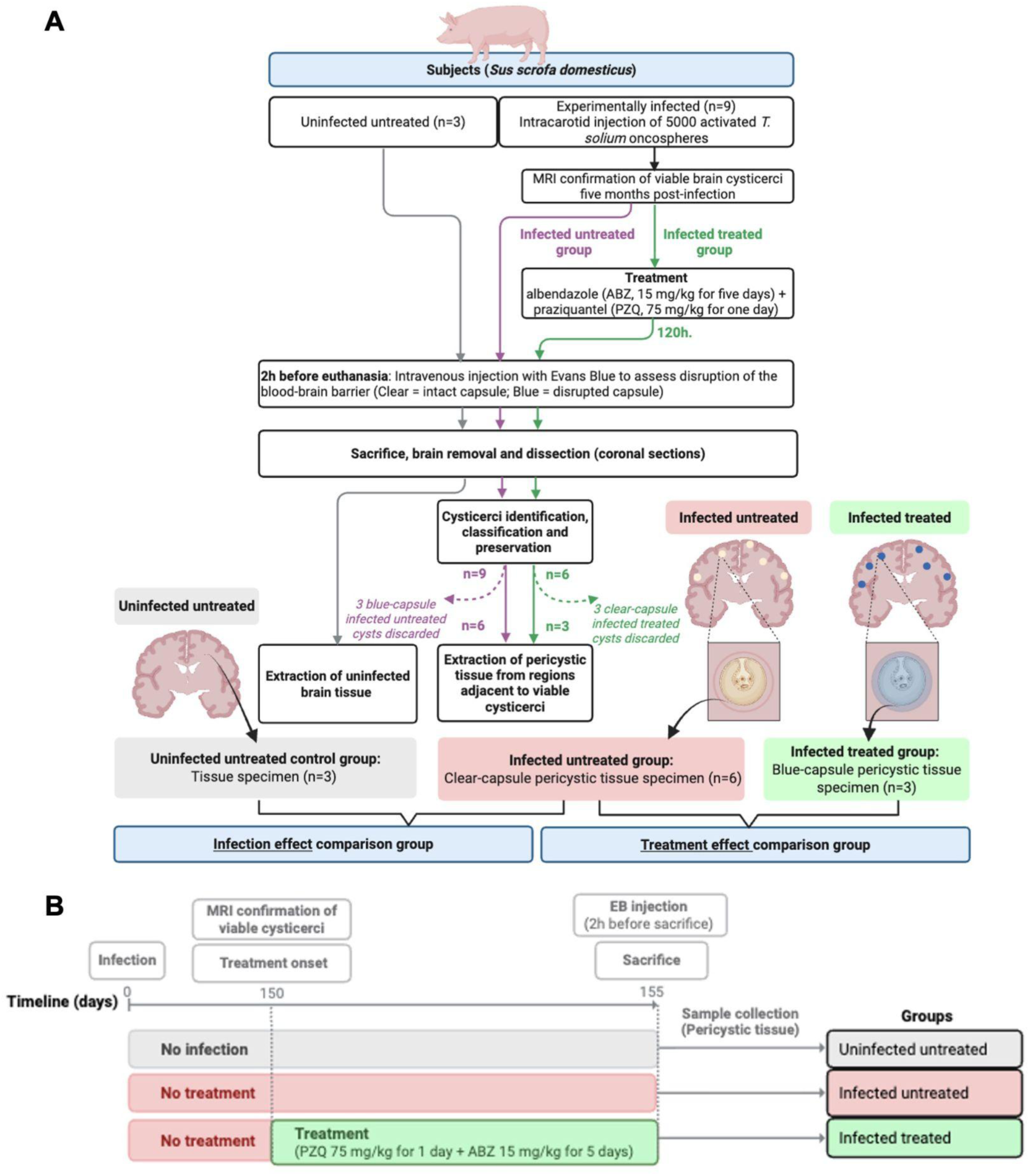
Experimental design and sample allocation. **(A)** Schematic overview of the three experimental groups, tissue selection strategy, and the two primary contrasts defined for differential expression analysis. All infected-untreated specimens derive from Clear-capsule cysts (intact BBB confirmed by Evans Blue); all infected-treated specimens derive from Blue-capsule cysts (BBB disruption confirmed by Evans Blue). **(B)** Infection and treatment timeline.

Two primary contrasts were defined. (i) First, to evaluate the effect of infection, we compared tissue from infected-untreated pigs with intact capsules to tissue from uninfected-untreated controls. (ii) Second, to assess the effect of antiparasitic treatment, we compared tissue from infected-treated pigs with disrupted capsules to tissue from infected-untreated pigs with intact capsules.

### RNA extraction and library preparation

Total RNA was extracted from corticomeningeal and parenchymal pericystic tissue using the QIAzol protocol followed by column purification. Samples were thawed on ice; tissue was removed from the preservation buffer using sterile forceps, trimmed to size, and the remainder returned to the buffer. Tissue was homogenised in 1 mL QIAzol using Macherey-Nagel Type G bead tubes in a TissueLyser (2 min at 25 Hz per side). RNA was isolated using the RNeasy Lipid Tissue Mini Kit (Qiagen, Hilden, Germany).

RNA quality was assessed by RNA ScreenTape on a TapeStation system (Agilent Technologies, CA, USA) and concentration quantified by Qubit 2.0 RNA HS Assay (Thermo Fisher Scientific, MA, USA). Only samples with DV200 ≥ 60% were retained. Ribosomal RNA was depleted with the Ribo-Zero Plus rRNA Depletion Kit (Illumina, CA, USA). Sequencing libraries were prepared using the NEBNext Ultra II Non-Directional RNA Library Prep Kit (New England BioLabs, MA, USA). Final library quality and fragment size distribution (mean ∼420 bp; insert size ∼300 bp) were confirmed by High Sensitivity D1000 ScreenTape. Libraries were multiplexed using Illumina 8-nt dual indices, pooled equimolarly, and sequenced on an Illumina NovaSeq platform in 150 bp paired-end mode targeting ∼40 million read pairs per sample.

### Quality assessment and filtering of raw data

All analyses were implemented in R v4.3.2 and Bash under Ubuntu 22.04; scripts and figure code are publicly available (see Data Availability). Raw FASTQ files were assessed with FastQC (v0.12) and MultiQC (v1.17). Adapter trimming and quality filtering were performed with BBDuk (BBTools v38): residual adapters were removed, read ends were trimmed to quality Q20, reads shorter than 35 bp after trimming were discarded, and low-complexity sequences were filtered.

### Transcript-level quantification

Transcript-level quantification used Salmon (v1.10) in quasi-mapping mode. A reference index was built from the *Sus scrofa domesticus* Sscrofa11.1 NCBI annotation using all annotated transcripts (k-mer length = 27). Trimmed paired-end reads were quantified using unstranded library mode, consistent with the non-directional library preparation protocol. Flags --validateMappings and --gcBias were enabled to improve mapping specificity and correct GC-content-related biases. Transcript-level count estimates, effective lengths, and TPM values were produced for downstream gene-level aggregation.

### Differential expression analysis

Quantification files were imported with tximport (v1.30) using length-aware transcript-to-gene aggregation. A tx2gene mapping table was derived from the Sscrofa11.1 GTF using rtracklayer (v1.62) and GenomicFeatures (v1.54). Genes with median count below 10 in fewer than three samples were excluded.

Then, differential expression analysis was performed with DESeq2 (v1.40), which models gene counts with a negative binomial generalised linear model and normalisation with the median-ratio method. Prior to model fitting, principal component analysis (PCA) of variance-stabilised expression data was performed to explore major sources of transcriptional variation across samples (Fig S1). Of the twelve samples included, eleven were derived from parenchymal tissue and one from corticomeningeal tissue. Because there was a single corticomeningeal specimen, tissue type could not be incorporated as a model covariate without consuming a degree of freedom that the dataset could not support; it was therefore excluded from the design. The absence of discernible clustering by tissue type in the PCA was consistent with this decision. The final model comprised the experimental group as the sole factor.

For each contrast, Wald tests provided log2 fold change (log2FC) estimates and associated p-values. To obtain accurate effect size estimates for reporting and visualisation, log2FC values were subsequently stabilised using the apeglm shrinkage estimator as implemented in the lfcShrink function of DESeq2 (v1.40). Genes with shrinkage-adjusted |log2FC| > 1 and Benjamini-Hochberg-adjusted p-value (FDR) < 0.05 were considered differentially expressed.

### Gene set enrichment analysis

Functional pathway enrichment used Gene Set Enrichment Analysis (GSEA) implemented in clusterProfiler (v4.10.1). Genes were ranked by the DESeq2 Wald statistic derived from the unshrunken results for each contrast; the unshrunken statistic was used for ranking because log2FC shrinkage can compress estimates for lowly expressed genes and distort gene-level rankings. KEGG pathway annotations provided the gene sets. Enrichment scores, normalised enrichment scores (NES), and FDR-adjusted p-values were computed for each contrast. Pathways with |NES| > 1 and FDR < 0.05 were considered significantly enriched.

## Results

### Viable NCC infection is associated with local immune activation and broad suppression of BBB-associated, vascular, and neuronal signalling programmes

To characterise the transcriptional impact of viable neurocysticercosis infection in the host, we first compared pericystic host brain tissue from infected-untreated pigs with intact viable lesions (n = 6) against uninfected brain tissue from uninfected-untreated controls (n = 3). Infection induced widespread gene expression changes, with 461 upregulated and 175 downregulated genes (**Supplementary Table 1; Fig. 2B**). Pathway enrichment analysis showed that these transcriptional changes were organised in four main biological axes: (i) immune activation, (ii) vascular, endothelial and BBB remodelling, (iii) neuroendocrine signalling, and (iv) neuronal activity (**Supplementary Table 2; Fig. 2C-F**).

**Fig 2.**
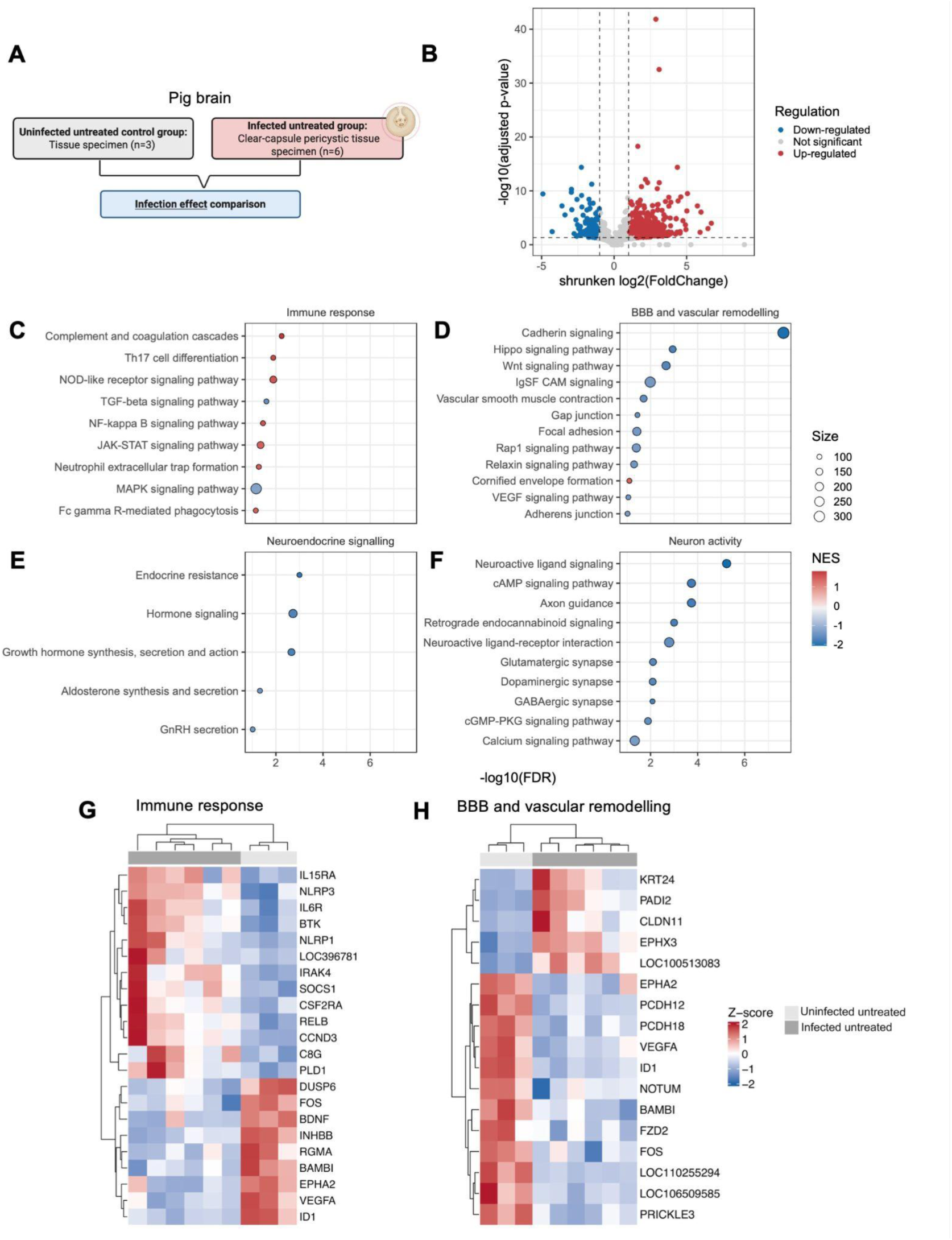
Transcriptional changes in pericystic brain tissue associated with viable NCC infection. **(A)** Study design for the infection-effect comparison: pericystic tissue from infected untreated pigs with Clear-capsule (viable) cysts vs brain tissue from uninfected untreated controls. **(B)** Volcano plot of differentially expressed genes; blue = downregulated, red = upregulated, grey = not significant; dashed lines indicate |log2FC| = 1 and FDR = 0.05 thresholds. **(C-F)** Enriched KEGG pathways in the **(C)** immune response, **(D)** BBB and vascular remodelling, **(E)** neuroendocrine signalling, and **(F)** neuronal activity categories. Dot colour represents NES (red = positive enrichment, blue = negative enrichment), dot size represents pathway gene-set size, and the x-axis shows -log10(FDR). **(G)** Heatmap of z-score-scaled regularised log expression for selected leading-edge immune-response genes. **(H)** Heatmap for BBB and vascular remodelling genes.

At the immune level, pericystic tissue showed increased expression of genes implicated in innate immune sensing and activation. Bruton’s tyrosine kinase (BTK) and the NF-κB family transcription factor RELB were among the most prominently upregulated genes (**Fig 2G**), alongside the phagosome-associated enzyme phospholipase D1 (PLD1). Consistent with this gene-level pattern, pathway enrichment analysis revealed positive enrichment of complement and coagulation cascades, Th17 cell differentiation, NOD-like receptor signalling, JAK-STAT signalling, and Fc gamma receptor-mediated phagocytosis (**Fig 2C**). In contrast, the MAPK signalling pathway was negatively enriched, a pattern accompanied by reduced expression of the MAPK-regulatory dual-specificity phosphatases DUSP6 and DUSP10, the immediate-early transcription factor FOS, and the neurotrophic factor BDNF. Genes associated with TGF-β signalling also showed coordinated downregulation.

Beyond these immune-related changes, viable infection was characterised by a reduction in expression of genes involved in BBB-associated, vascular, and endothelial programmes (**Fig 2H**). Angiogenic and endothelial regulators, including VEGFA, ID1, DLL4, EPHA2, and FZD2, were all downregulated in pericystic tissue. Junctional and adhesion genes were also affected, as the protocadherins PCDH12 and PCDH18 showed reduced expression, while the tight-junction component CLDN11 was upregulated, suggesting selective remodelling of junctional composition rather than a uniform collapse of barrier function. At the pathway level, Cadherin signalling, Hippo signalling, Wnt signalling, IgSF CAM signalling, VEGF signalling, vascular smooth muscle contraction, adherens junction, and gap junction were all negatively enriched (**Fig 2D**), reinforcing a broad suppression of vascular and endothelial programmes in the pericystic space.

Having identified changes in gene expression of immune and BBB remodelling pathways during viable infection, we next examined whether these alterations extended to other processes. Infection was additionally associated with negative enrichment of neuroendocrine signalling pathways, including growth hormone synthesis, secretion and action; aldosterone synthesis and secretion; and GnRH secretion (**Fig 2E**). Pathways governing neuronal activity were similarly suppressed, including neuroactive ligand-receptor interaction, glutamatergic, GABAergic, and dopaminergic synapses, and cAMP, cGMP-PKG, and calcium signalling (**Fig 2F**). At the gene level, OPRM1, HOMER1, TRPC3, and RGMA, all associated with synaptic function and neuronal plasticity, showed reduced expression in infected pericystic tissue (**Supplementary Table 1**).

Taken together, the transcriptional profile of pericystic tissue during viable NCC infection combines evidence of localised innate immune engagement with broad suppression of vascular, BBB-associated, and neuronal signalling programmes. These findings are consistent with a pericystic environment that is immunologically active yet neurovascularly quiescent around intact, viable cysts, though experimental validation is required before causal interpretation is warranted.

### Post-treatment BBB disruption is associated with inflammatory signalling and a directional shift in BBB-associated and vascular gene programmes

To characterise the pericystic transcriptional environment following antiparasitic treatment with confirmed BBB disruption, we compared infected treated samples with infected untreated samples. Although treatment induced fewer transcriptional changes than infection, with 160 upregulated and 57 downregulated genes (**Supplementary Table 3; Fig. 3B**), pathway enrichment analysis showed that these changes converged on three biological axes also observed in the infection response (i) immune activation and (ii) BBB/tissue remodelling and (iii) neuroendocrine signalling. However, in treatment, these changes were additionally accompanied by pathways related to controlled cell death (**Supplementary Table 4; Fig. 3C-F**). Given that the treated group comprised only three animals, these findings represent preliminary observations and should be interpreted with appropriate caution.

**Fig 3.**
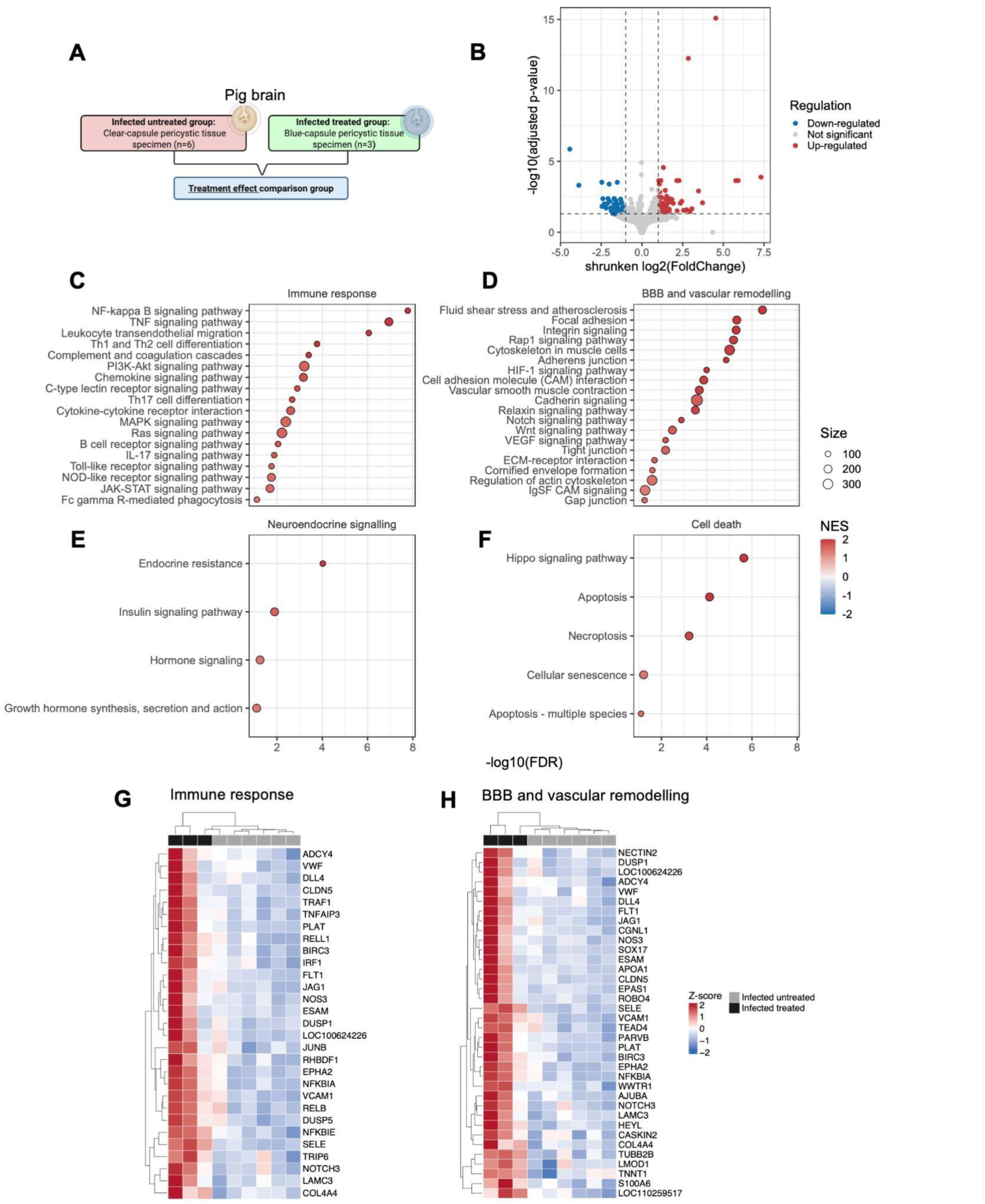
Transcriptional changes in pericystic brain tissue associated with antiparasitic treatment. **(A)** Study design for the treatment-effect comparison: pericystic tissue from infected treated pigs with Blue-capsule (disrupted) cysts vs pericystic tissue from infected untreated pigs with Clear-capsule (viable) cysts. **(B)** Volcano plot of differentially expressed genes; colours and thresholds as in Fig 2B. **(C-F)** Enriched KEGG pathways in the **(C)** immune response, **(D)** BBB and vascular remodelling, **(E)** neuroendocrine signalling, and **(F)** cell death categories. Dot colour, size, and x-axis as in Fig 2C-F. **(G)** Heatmap for selected immune-associated genes. **(H)** Heatmap for BBB, vascular, and endothelial remodelling genes.

The immune transcriptional response in treated tissue was markedly broader than that observed in infected untreated tissue. GSEA revealed robust positive enrichment of NF-κB signalling, TNF signalling, leukocyte transendothelial migration, Th1 and Th2 cell differentiation, complement and coagulation cascades, PI3K-Akt signalling, chemokine signalling, cytokine-cytokine receptor interaction, MAPK signalling, Toll-like receptor signalling, NOD-like receptor signalling, and JAK-STAT signalling (**Fig 3C**). At the gene level, this pattern was driven in part by increased expression of NF-κB regulatory feedback components NFKBIA, NFKBIE, and TNFAIP3, as well as the adaptor TRAF1 and the transcriptional regulators RELB, IRF1, and JUNB (**Fig 3G**). The simultaneous induction of both upstream NF-κB activators and their negative feedback regulators suggests an acute, self-limiting inflammatory response consistent with the release of parasite antigens into the pericystic environment following cyst capsule disruption.

The most distinctive feature of the treatment-associated transcriptional profile was a pronounced increase in expression of genes involved in endothelial activation, vascular regulation, and tissue remodelling (**Fig 3D**). These included adhesion molecules such as VCAM1 and SELE, vascular-associated genes including VWF, PLAT and EPHA2, and endothelial junction components such as CLDN5, ESAM, and NECTIN2 (**Fig 3H**). Genes involved in extracellular matrix structure, including COL4A4 and LAMC3, were also upregulated. Furthermore, treatment was associated with increased gene expression of several Notch-associated components, including ligands DLL4 and JAG1 in addition to NOTCH3 receptor, suggesting that the post-treatment vascular response may also involve Notch-related endothelial or neurovascular signalling. Additional upregulated genes in treated samples included FLT1, NOS3, PARVB, SEMA3G, ADCY4, and DUSP1, all of which participate in vascular and endothelial signalling.

Treatment was additionally associated with positive enrichment of cell-death-related pathways, including Hippo signalling, apoptosis, necroptosis, and cellular senescence (**Fig 3F**), indicating that disrupted cyst tissue harbours concurrent inflammatory, endothelial-activation, and structural-stress transcriptional programmes.

Overall, these findings suggest that antiparasitic treatment shifts the pericystic transcriptional environment towards a state of heightened inflammation, endothelial activation, and vascular remodelling, a pattern consistent with the host response to cyst capsule disruption and the release of parasite-derived material into the pericystic space.

### Integrated analysis shows shared immune engagement and opposing neurovascular programmes between infection and treatment

To compare the transcriptional programmes associated with viable infection and antiparasitic treatment, we integrated the pathway enrichment results from both contrasts using paired NES values (**Fig 4A**), examined the gene-level overlap between conditions as an UpSet plot (**Fig 4B**), and visualised the expression patterns of shared differentially expressed genes across all three experimental groups (**Fig 4C**).

**Fig 4.**
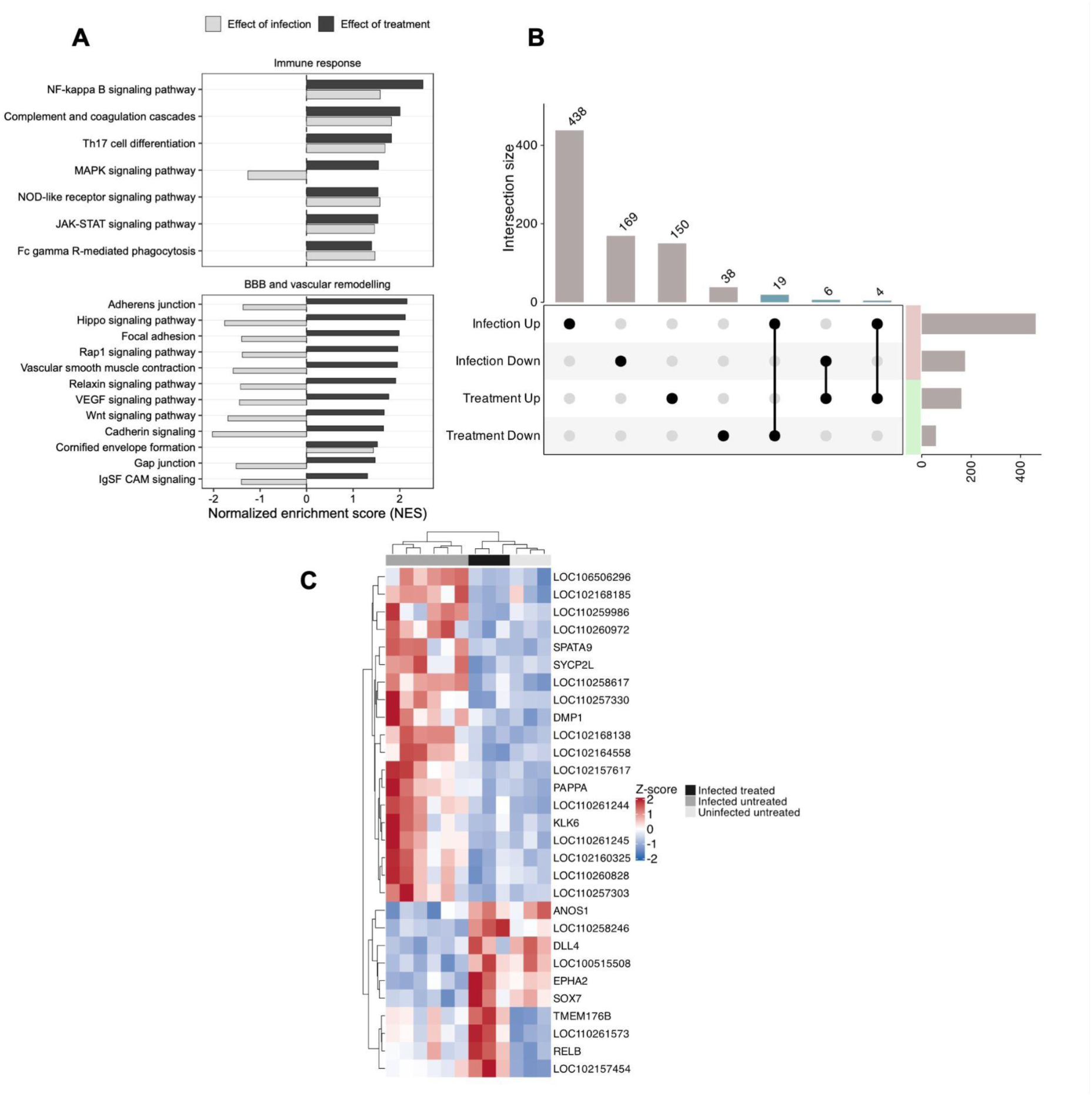
Integration of transcriptional programmes across viable infection and post-treatment pericystic tissue. **(A)** Paired bar chart comparing NES values for selected KEGG pathways in the infection-effect (light grey) and treatment-effect (black) contrasts, grouped into immune response (upper panel) and BBB and vascular remodelling (lower panel). Positive bars indicate positive enrichment; negative bars indicate negative enrichment. **(B)** UpSet plot of differentially expressed genes unique to and shared across infection upregulation, infection downregulation, treatment upregulation, and treatment downregulation. Intersection sizes are labelled above bars; set sizes are shown to the right. **(C)** Heatmap of z-score-scaled regularised log expression for selected shared differentially-expressed genes across all three experimental groups.

A core set of immune pathways was positively enriched under both conditions. NF-κB signalling, complement and coagulation cascades, Th17 cell differentiation, NOD-like receptor signalling, JAK-STAT signalling, and Fc gamma receptor-mediated phagocytosis showed concordant positive NES in both the infection and treatment contrasts (**Fig 4A**), indicating that local immune engagement is a persistent feature of pericystic tissue that precedes and extends beyond cyst disruption. In contrast, MAPK signalling showed opposing enrichment, negative during viable infection and positive after treatment, the most pronounced directional divergence in the immune panel. All BBB-associated and vascular pathways followed the same opposing logic: negatively enriched during infection and positively enriched after treatment (**Fig 4A**), reinforcing the directional reversal of the neurovascular axis between the two disease states.

Despite the shared immune axis, the gene-level overlap between conditions was limited (**Fig 4B**). The large majority of DEGs were condition-specific: 438 were uniquely upregulated in infection, 169 uniquely downregulated in infection, and 150 uniquely upregulated in treatment. Only 19 genes were upregulated in both conditions, 6 downregulated in both, and 4 showed discordant regulation. The heatmap of shared DEGs (**Fig 4C**) captured this separation visually, revealing three expression clusters. The first cluster showed increased expression after infection but reduced after treatment. The annotated genes in this cluster included PAPPA, encoding a metalloproteinase involved in extracellular matrix remodelling and IGF signalling; KLK6, a serine protease associated with tissue remodelling; and ANOS1, involved in extracellular matrix organisation. The second cluster, containing the Notch ligand DLL4, the receptor tyrosine kinase EPHA2, and SOX7, a transcription factor implicated in endothelial specification, showed reduced expression during infection but marked upregulation in treated animals, reinforcing the treatment-specific shift towards endothelial activation identified in the previous analysis. The third cluster, which included TMEM176B, an inflammasome-associated transmembrane protein, and RELB, a transcriptional regulator of the non-canonical NF-κB pathway, was upregulated in both infection and treatment, consistent with sustained innate immune engagement across viable infection and post-treatment states.

These integrated findings indicate that viable NCC infection and early post-treatment cyst disruption share a persistent immune engagement axis but diverge markedly in their BBB-associated, vascular, and MAPK-related transcriptional programmes. The directional reversal of the neurovascular axis, combined with the largely non-overlapping gene-level responses, constitutes the principal transcriptional signal emerging from this first survey of pericystic brain tissue in NCC.

## Discussion

Neurocysticercosis pathology is fundamentally shaped by parasite viability and the immunological microenvironment of the pericystic space, yet the molecular programmes underlying these states have remained undefined at the whole-transcriptome level. Here we address this knowledge gap by presenting the first bulk RNA-seq analysis of the pericystic brain during *T. solium* infection and subsequent antiparasitic treatment using a clinically relevant porcine model. Three principal patterns emerge from this study. First, local immune engagement is a shared feature of both viable-infection and post-treatment tissue; second, MAPK signalling shows divergent enrichment between conditions; and, finally, BBB-associated and vascular remodelling programmes undergo the most pronounced directional reversal, suppressed during viable infection and activated following treatment. Because this is an exploratory, first-in-model dataset with a small treated group (n = 3) and no orthogonal protein-level validation, all findings are presented as hypothesis-generating observations.

First, local immune engagement was evident in both viable infection and post-treatment tissue, with enrichment of inflammatory programmes including NF-κB, NOD-like receptor, JAK-STAT, Fc gamma receptor-mediated phagocytosis, and complement/coagulation cascades. The induction of NF-κB-responsive feedback genes such as NFKBIA, NFKBIE, and TNFAIP3 supports the presence of a well-described active but regulated inflammatory signalling in cyst-associated tissue (5). The upregulation of BTK during infection is notable because BTK acts upstream of several innate immune signalling cascades, including TLR-NF-κB and NLRP3-related inflammatory responses in myeloid cells (14); however, the current data are limited to mRNA expression and provide no functional or protein-level evidence. BTK is noted here solely as a candidate inflammatory node for future mechanistic investigation in NCC. It is also worth noting that transcriptional upregulation of immune-associated genes does not necessarily imply equivalent protein-level inflammatory activity. *T. solium* cysticerci have been shown to deploy post-transcriptional immune evasion mechanisms, including degradation and molecular mimicry of host immunoglobulins, that may modulate the functional output of host immune gene expression (4). Whether the transcriptional immune signatures identified here translate to corresponding protein-level inflammatory responses, or are partially attenuated by parasite-derived post-transcriptional mechanisms, cannot be determined from the present bulk RNA-seq data and warrants investigation at the protein and functional levels.

A more distinctive finding was the divergent behaviour of genes participating in MAPK signalling between viable infection and treatment. In particular, such changes might be relevant to the epileptogenic potential of NCC. This disease is widely recognised as a major cause of acquired epilepsy in endemic regions, and has been proposed as a natural model of human epileptogenesis (15). In our data, MAPK signalling showed opposite enrichment patterns between viable infection and treatment, accompanied by contrasting expression of genes involved in MAPK regulation and neuronal activity. Specifically, DUSP6, DUSP10, FOS, and BDNF were downregulated in infected tissue, whereas DUSP1 and DUSP5 were upregulated in treated pericystic tissue. This pattern may provide insight into the regulatory mechanisms underlying the differences in MAPK-associated pathway enrichment between conditions, since DUSPs are important regulators of MAPK signalling and BDNF has been shown to influence DUSP1 expression in neural cells (16). The biological relevance of these changes is supported by the established roles of BDNF/TrkB signalling, MAPK/ERK pathways, and immediate-early genes such as FOS in neuronal plasticity, excitability, and epileptogenic processes (17). In particular, reduced BDNF expression in the viable infection state may suggest altered neuronal support signalling in pericystic tissue, since BDNF is known to contribute to maintaining neuronal integrity and synaptic plasticity (18). BDNF is also measurable in peripheral blood and has been explored as a biomarker in epilepsy and other neurological disorders, although its interpretation is highly context-dependent (19). Therefore, reduced BDNF expression may be most useful as an exploratory signal of altered neuronal support and seizure-relevant plasticity pathways. Critically, this study does not measure seizure activity or neuronal excitability; these findings represent candidate signals warranting dedicated mechanistic investigation rather than conclusions about seizure susceptibility.

Finally, the changes associated with BBB remodelling suggest that antiparasitic treatment is accompanied by endothelial activation and vascular remodelling. In our analysis, infection and treatment showed opposing patterns in vascular, endothelial, and BBB-related pathways. These programmes were relatively suppressed during viable infection but increased after treatment, consistent with previous NCC studies linking BBB disruption and vascular changes to inflammatory lesion progression (13,20). At the gene level, this pattern was supported by increased expression of endothelial activation and leukocyte adhesion markers, including VCAM1, SELE, ICAM1, and VWF. Among these, soluble VCAM1 has been linked to impaired brain endothelial barrier function, suggesting its potential value as an accessible indicator of vascular activation after treatment onset (21). Beyond adhesion markers, treatment also altered vascular signalling pathways which may help explain this endothelial activation state. First, the observed pattern was accompanied by Notch-associated changes: DLL4 was downregulated during viable infection but upregulated after treatment, while NOTCH3 and JAG1 were upregulated only in the post-treatment state. Given its role in vascular regulation, BBB permeability, and inflammatory infiltration, this pattern may implicate Notch-associated signalling in the post-treatment lesion response (22–24). Second, the VEGF-related signal appeared to reflect pathway remodelling rather than uniform VEGF activation. VEGFA was downregulated in both conditions, whereas receptor and downstream components, including FLT1 and NOS3, were increased after treatment. The upregulation of NOS3, encoding endothelial nitric oxide synthase, may reflect activation of endothelial vasodilatory and leukocyte-adhesion signalling pathways (25), though NOS3 can also contribute to oxidative tissue injury in inflammatory context (26). Overall, these findings suggest that post-treatment disrupted lesions acquire a transcriptional profile compatible with endothelial activation, leukocyte trafficking and vascular remodelling, while identifying Notch-associated and VEGF/NOS signalling as candidate pathways for future mechanistic evaluation. It should be noted that the treatment-effect contrast in this study captures the combined biological state of post-treatment cyst disruption and confirmed BBB disruption, and cannot dissociate the transcriptional contributions of the antiparasitic drugs themselves from those arising from cyst capsule breakdown or barrier disruption *per se.* This confound is inherent to the study design and to the biology of NCC treatment response, in which BBB disruption and cyst disruption co-occur as features of the same pathological state. Future studies incorporating animals treated but without BBB disruption, or longitudinal sampling before and after treatment onset, would be required to isolate these contributions.

Taken together, these findings have several implications for understanding neurocysticercosis pathogenesis. They provide evidence of local immune engagement alongside altered neuronal-support signalling and vascular programmes at the molecular level. More broadly, these transcriptional signatures provide candidate pathways of interest that include MAPK-related genes, VCAM1, Notch-associated signalling, and endothelial nitric oxide signalling. This could guide future studies of lesion activity, treatment response, and accessible biomarkers of neurovascular injury in NCC. However, the principal limitations of this study must be also acknowledged explicitly. The treated group comprised only three animals, which intrinsically limits statistical power despite the suitability of DESeq2’s negative binomial modelling framework for low-replicate designs. The treatment-effect transcriptional signatures should accordingly be considered preliminary hypotheses rather than definitive findings, and require validation in larger cohorts with a prospective power calculation. All findings are based on bulk RNA-seq without orthogonal confirmation by RT-qPCR, immunohistochemistry, or protein-level assays; the gene-level associations reported, including VCAM1, DLL4, NFKBIA, and BDNF, require independent experimental confirmation before clinical or therapeutic inference is warranted. The cross-sectional design, necessitated by the terminal nature of brain tissue sampling, precludes within-cyst longitudinal inference and limits causal interpretation of the observed transcriptional transitions. Finally, bulk RNA-seq cannot resolve the cellular sources of the identified transcriptional signatures; future spatially resolved and single-cell profiling studies will be essential to determine which cell populations drive the immune, endothelial, and neuronal changes observed in pericystic tissue across lesion states.

Ultimately, this study provides the first whole-transcriptome characterisation of pericystic brain tissue in a porcine model of neurocysticercosis, revealing distinct and partially opposing transcriptional states associated with viable parasite persistence and early post-treatment cyst disruption. Tissue surrounding intact viable cysts displays gene expression of localised immune engagement alongside broad suppression of BBB-associated molecular signatures. Following antiparasitic treatment, disrupted lesions acquire a markedly different transcriptional identity, characterised by intensified inflammatory signalling, endothelial activation and neurovascular remodelling. Together, these findings establish that NCC pathology involves coordinated and state-specific immune and neurovascular responses at the molecular level, and provide a candidate transcriptomic framework to guide future studies aimed at defining the cellular drivers of lesion transitions, identifying accessible biomarkers of lesion activity and treatment response, and evaluating adjunctive therapeutic strategies targeting the neurovascular and inflammatory axes in NCC.

## Statements

### Data Availability Statement

The complete bioinformatics analysis pipeline and figure-generation code are publicly available at https://github.com/carlaapaza/PigBrain2026. Raw Illumina sequencing reads have been deposited in the NCBI Gene Expression Omnibus (GEO) under accession number GSE334289.

### Author contributions

Conceptualization: CAAQ, SAGG, HHG, MZ. Data curation: CAAQ. Formal analysis: CAAQ, CCRP, EKPN. Funding acquisition: HHG. Investigation: CAAQ, SAGG, JAB, GA, HHG. Methodology: CAAQ, HHG, MZ. Visualization: CAAQ. Supervision: HHG, MZ. Writing - original draft: CAAQ, CCRP, EKPN, GA. Writing - review & editing: SAGG, JAB, GA, RHG, HHG, MZ.

## Supplementary information

**Supplementary Table 1. DESeq2 differential expression results for the infection effect.** Complete gene-level output from the comparison between infected untreated and uninfected control samples, including log2 fold changes, test statistics, nominal p-values and adjusted p-values.

**Supplementary Table 2. KEGG pathways significantly enriched for the infection effect.** Gene set enrichment analysis results for the infection-effect comparison, including pathways with absolute normalised enrichment score (|NES|) > 1 and false discovery rate (FDR) < 0.05.

**Supplementary Table 3. DESeq2 differential expression results for the treatment effect.** Complete gene-level output from the comparison between treated infected and untreated infected samples, including log2 fold changes, test statistics, nominal p-values and adjusted p-values.

**Supplementary Table 4. KEGG pathways significantly enriched for the treatment effect.** Gene set enrichment analysis results for the treatment-effect comparison, including pathways with absolute normalised enrichment score (|NES|) > 1 and false discovery rate (FDR) < 0.05.

**Figure S1. Principal component analysis of samples.** PCA plot based on variance-stabilized gene expression data, showing the distribution of samples based on **(a)** experimental group and **(b)** location of the sample.

