## Supplementary figures and images for "Pericystic brain transcriptomics reveals molecular signatures of immune activation and neurovascular remodelling in viable and post-treatment porcine neurocysticercosis"

### Fig S1.tiff

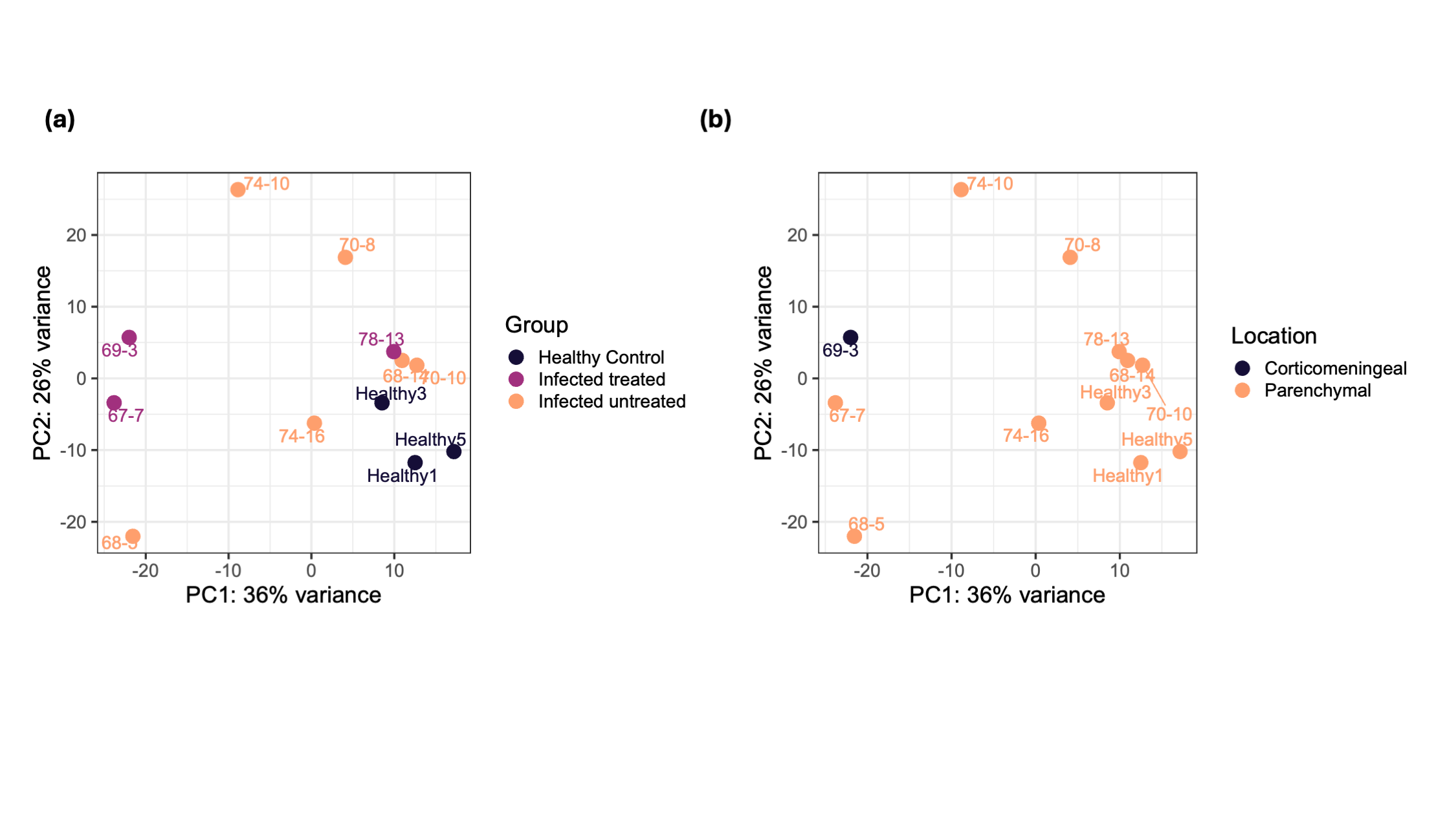
